# The blobulator: a toolkit for identification and visual exploration of hydrophobic modularity in protein sequences

**DOI:** 10.1101/2024.01.15.575761

**Authors:** Connor Pitman, Ezry Santiago-McRae, Ruchi Lohia, Ryan Lamb, Kaitlin Bassi, Lindsey Riggs, Thomas T. Joseph, Matthew E.B. Hansen, Grace Brannigan

## Abstract

While contiguous subsequences of hydrophobic residues are essential to protein structure and function, as in the hydrophobic core and transmembrane regions, there are no current bioinformatics tools for module identification focused on hydrophobicity. To fill this gap, we created the *blobulator* toolkit for detecting, visualizing, and characterizing hydrophobic modules in protein sequences. This toolkit uses our previously developed algorithm, blobulation, which was critical in both interpreting intra-protein contacts in a series of intrinsically disordered protein simulations [1] and defining the “local context” around disease-associated mutations across the human proteome [2]. The *blobulator* toolkit provides accessible, interactive, and scalable implementations of blobulation. These are available via a webtool, a VMD plugin, and a command line interface. We highlight use cases for visualization, interaction analysis, and modular annotation through three example applications: a globular protein, two orthologous membrane proteins, and an IDP. The *blobulator* webtool can be found at www.blobulator.branniganlab.org, and the source code with pip installable command line tool, as well as the VMD plugin with installation instructions, can be found on GitHub at www.GitHub.com/BranniganLab/blobulator.

## 1 INTRODUCTION

Protein sequences are modular: they consist of contiguous units (such as alpha helices, functional domains, etc.) which are incorporated into a range of analysis pipelines, visualization tools, and conceptual frameworks. Few tools for detecting modularity, however, explicitly incorporate residue hydrophobicity, despite the cooperative nature of the hydrophobic effect; the critical role of hydrophobicity in stabilizing the core of globular proteins; and the commonplace use of residue hydrophobicity in predictors of protein disorder [3–8] and membrane interactions [9, 10]. We introduced “blobulation” as a scheme for segmenting protein sequences by contiguous hydrophobicity, motivated by the need for an analog to secondary structure when reducing the dimensionality of intrinsically disordered protein (IDP) simulations [1]. Although blobulation was first applied to intrinsically disordered proteins, contiguously hydrophobic regions are most frequently found in the buried cores of structured proteins and have functional and evolutionary signatures in the human proteome: disease-associated single-nucleotide polymorphisms (dSNPs) are more likely to be found in contiguously hydrophobic regions and are particularly likely to bridge two such regions [2]. These results support the interpretation of hydrophobic blobs as evolutionarily constrained interaction “nodes” within a protein sequence. Here, we introduce a toolkit designed to make blobulation accessible and convenient for a wide range of users.

Blobulation detects modularity in protein sequences by searching for “h-blobs”: subsequences longer than a threshold length in which the predicted hydrophobicity score for every residue (its “hydropathy”) exceeds a user-provided threshold. The remaining subsequences are sorted into “p-blobs” (which satisfy the length, but not the hydropathy criterion) and “s-blobs” (which satisfy neither length nor hydropathy criteria). While secondary structure detectors typically use a default set of parameters that users rarely change, blobulation settings are meant to be tuned: adjusting the hydropathy and length thresholds in blobulation allows users to gradually shift from detecting many small modules to a few longer ones, bringing the relevant aspects of sequence organization into focus (use-case examples shown in section 4). Furthermore, since blobulation requires only the sequence, it is particularly appropriate in scenarios where circumventing structure is desirable (including IDPs).

Many existing tools cluster residues into protein modules using secondary structure. While blobulation yields modules analogous to secondary structure elements, we have not previously provided “blob” versions of tools that traditionally rely on secondary structure. On the left half of Table 1, we list the common use-cases for secondary structure identification among currently available tools: residue characterization (secondary structure of each residue), module identification (determining where an alpha helix, beta sheet, or other secondary structure element begins and ends), and module characterization (calculating collective properties of the secondary structure element, for instance, the net charge per residue within a helix or beta sheet). The columns show the properties used to define modularity (secondary structure or hydropathy), and then are subdivided by the functionality of the tools: whether the tool makes modularity predictions (usually via a command-line interface), or provides a graphical interface (via a sequence or structure view). For secondary structure tools, module prediction can be accessed through command-line tools [12, 13, 28], graphically displayed alongside the sequence [14–18], or incorporated into visualizations of the structure [19–21]. As shown on the right half of Table 1, tools for characterization [22], annotation [23–26], or coloring by the hydropathy of individual residues [19, 20, 27] are well-established. Yet, few tools incorporate “modular” definitions based on contiguous hydrophobicity, and we are unaware of any that allow graphical exploration or module characterization. Additionally, while tools that provide plots of residue hydropathy may invite the viewer to identify contiguous hydrophobic regions by eye, they do not provide a systematic, automatable version of the same process.

**Table 1.** Features of the *blobulator* toolkit compared to other available tools for characterizing protein elements by secondary structure or hydropathy. Rows vary by the elements characterized or identified, and columns vary by the type of analysis or visualization. Each cell gives example of tools available for this purpose as well as the target users of the most accessible tool. Features that can be accessed via a GUI are considered accessible to end-users, while features that require significant scripting are noted as requiring power users. Blobulation completes in under a second for typical proteins and scales more efficiently with sequence length than a widely-used secondary structure predictor (Figure S6).

|  | Secondary Structure |  |  | Hydropathy |  |  |
| --- | --- | --- | --- | --- | --- | --- |
|  | Prediction | Sequence View | Structure View | Prediction | Sequence View | Structure View |
| **Residue Characterization<sup>a</sup>** | end user<br>(e.g. S4Pred [11], SCRATCH [12], AlphaFold [13]) | end user<br>(e.g. PROTEUS2 [14], PSIPRED [15], JPred4 [16], NetSurfP [17], ALIGNSEC [18]) | end user<br>(e.g. VMD [19], UCSF Chimera [20], POLYVIEW [21]) | end user<br>(e.g. EMBL-EBI [22]) | end user<br>(e.g. BioJava [23], Ex-pasy [24], MPEx [25], EMBOSS [26]) | end user<br>(e.g. VMD [19], UCSF Chimera [20], UGENE [27]) |
| **Module Identification<sup>b</sup>** |  |  |  | power user<br>(**blobulator** CLI) | end user<br>(**blobulator** webtool) | end user<br>(**blobulator** VMD) |
| **Module Characterization<sup>c</sup>** | power user<br>(e.g. S4Pred [11], SCRATCH [12], AlphaFold [13]) | (not widely available) | power user<br>(e.g. VMD [19], UCSF Chimera [20]) |  |  |  |

<sup>a</sup> predicts and/or visualizes secondary structure or hydropathy by residue
<sup>b</sup> predicts and/or visualizes stretches of contiguous secondary structure or hydropathy
<sup>c</sup> predicts and/or visualizes physicochemical or other properties aggregated over the topological element

To provide a previously unavailable analogue to secondary structure tools that defines and characterizes modules based on hydropathy (bottom right of Table 1), we developed the *blobulator* toolkit: an accessible, interactive, and scalable suite of tools that includes a webtool, command line interface (CLI), and a Visual Molecular Dynamics (VMD) [19] plugin. The webtool supports a sequence-level view of blobs that includes smooth real-time parameter adjustment, a feature to introduce mutations, and layers of annotations including physicochemical properties and disease-associated mutations. The backend program for the webtool is available as a pip-installable CLI and supports high-throughput batch blobulation of FASTA files. Finally, the VMD plugin allows users to view blobs on protein structures by creating a blobulator interface in VMD, which stores identified blobs for analysis across simulation trajectories and the generation of publication-quality images and movies. As a whole, the *blobulator* toolkit provides previously unavailable functionality for detecting modules defined by hydrophobicity through both prediction and visualization of blobs and their characteristics in protein sequence and structure. Additionally, it goes beyond secondary structure analogues, enabling end users to access module characterization through graphical user interfaces.

In this paper, we outline the blobulation algorithm and each component of the *blobulator* toolkit. We then demonstrate its utility in three example applications: detecting blobs corresponding to tertiary interactions in a globular protein; comparing blobulated sequences from a structurally conserved membrane protein family; and quantifying the effects of disease-associated mutations in an IDP. We close by describing future applications for these tools, including novel insights that the *blobulator* is positioned to enable.

## 2 BLOBULATION

Whole-sequence blobulation consists of two steps: digitization and clustering. We provide a description of the algorithm below, as well as a pseudocode version of the algorithm in the SI (Supplementary Methods).

1. In the digitization step, the algorithm defines a given residue as hydrophobic or non-hydrophobic as follows:

a. The user selects a normalized (0 to 1) hydropathy scale, and sets a threshold hydropathy (*H*^∗^) on this scale.
b. The hydropathy for each residue (*i*) is assigned based on the selected normalized hydropathy scale. The hydropathies are then smoothed for each residue *i* and the residues adjacent to it in sequence (*i* − 1, *i*, and *i* + 1) , yielding the smoothed hydropathy *H_i_*.
c. Residue *i* is classified as hydrophobic (*H_i_ > H*^∗^) or non-hydrophobic (*H_i_* ≤ *H*^∗^), shown in Fig. 1A.
2. In the clustering step, subsequences termed “blobs” are defined as follows:

a. The user sets *L*_min_ equal to a positive integer representing a threshold number of residues required to form an h- or p-blob.
b. Subsequences of at least *L*_min_ sequential hydrophobic residues are classified as hydrophobic blobs (h-blobs) (Fig. 1B).
c. All other linking sequences are then classified based on their *L*_min_ as either non-hydrophobic blobs (p-blobs, *L* ≥ *L*_min_) or short blobs (s-blobs, *L < L*_min_) (Fig. 1B).

**Figure 1.**
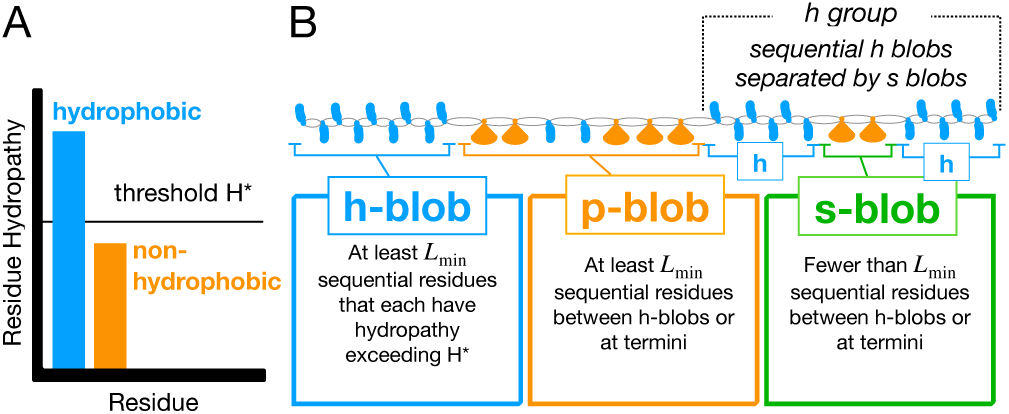
Blobulation algorithm. A) First, the sequence is digitized. Residues are classified as either hydrophobic (blue) or non-hydrophobic (orange) by comparing their hydropathy to the user-selected threshold, *H*^∗^. B) The sequence is then segmented into h-blobs, s-blobs, and p-blobs based on *H*^∗^ and *L*_min_. Used with permission from [2], Copyright 2022 PNAS.

Though there are default settings for the initial blobulation, the user can tune various parameters. Different combinations of *H*^∗^ and *L*_min_ will detect blobs with varying properties, and adjusting them can extract blobs that correspond to key hydrophobic regions (some examples are shown in section 4). Alternatively, users may reveal hierarchical layers of organization by blobulating a given sequence under varying parameter settings. Very high thresholds (*H*^∗^ approaching 1) will return one large p-blob, and very low thresholds (*H*^∗^ approaching 0) will return one large h-blob, while intermediate thresholds will reveal intrinsic segmentation within the protein sequence. Additionally, high *L*_min_ will only yield h-blobs if used with a low to moderate *H*^∗^.

We will also consider higher-order organization beyond individual blobs: “blob groups” are h-blobs separated only by s-blobs. Examples and further discussion of blob groups can be found in section 4. Blobs are labeled as follows: by their type (h, p, s), group number (1, 2, 3), and for h-blobs within a blob group, by a sub-group letter (a, b, c). This forms a unique label for every blob (h1a, s1, h1b, etc). We refer to the distribution of blobs in a protein sequence as its “blob topology”.

## 3 THE *BLOBULATOR* TOOLKIT

Here, we present three tools by which an amino acid sequence can be blobulated: a webtool, a command line interface (CLI), and a Visual Molecular Dynamics (VMD) plugin. As shown in the schematic in Figure S1, all three tools share the core blobulation algorithm outlined in section 2. The webtool can be used to interactively explore blobulated sequences, introduce mutations, and investigate various blob properties. The command line interface is the backend for the webtool, but also can be used to run batch processes and accepts DNA sequences. The VMD plugin can be used with molecular structures for analysis and creating images and movies of blobulated proteins in VMD. All images in this paper reflect the output of the *blobulator* version 1.0b.

### 3.1 Webtool

Blobulation of any amino acid sequence can be achieved using the *blobulator* webtool found at www.blobulator.branniganlab.org. On the “New Query” tab of the homepage, the user can submit either a database ID or manually enter a protein sequence for blobulation. From here, a user can blobulate any protein or amino acid sequence using an accepted ID or manually entering the sequence directly, as well as toggle to other information tabs. Figure 2 illustrates the blobulation of insulin, a peptide hormone that promotes glucose absorption by cells.

**Figure 2.**
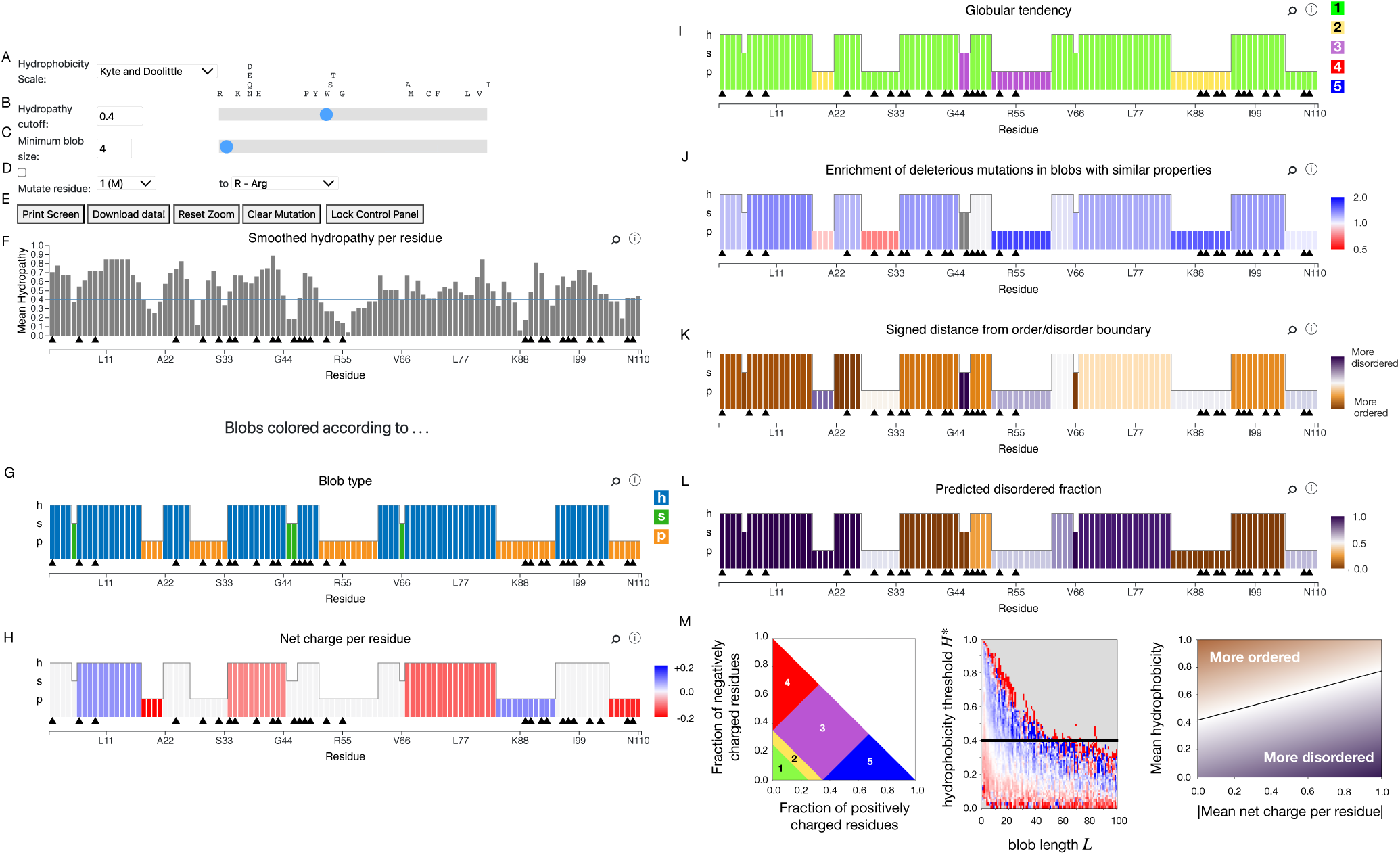
Screenshots from the *blobulator* webtool results page. Parameter and sequence adjustments are made via: a dropdown menu for selecting a hydropathy scale (A), hydropathy cutoff *H*^∗^ adjustment (numerical entry, slider, and 1-letter amino acid code) (B), and minimum h- or p-blob length adjustment *L*_min_ (numerical entry and slider) (C). Introducing mutations is done using the “mutate residue” panel (D). Additional data and display options: print screen, download data, reset zoom, clear mutation, and lock control panel (E). The “smoothed hydropathy per residue” track displays both residue hydropathy (gray bars) and *H*^∗^ (blue, horizontal line) (F). Clickable black triangles introduce known disease-associated mutations (only available via ID entry of a human sequence), and zooming is supported on all tracks. Blob type is represented by both bar height (y-axis) and color on the “blob type” track (height is preserved in subsequent tracks) (G). Blobs are then colored according physicochemical properties: H) net charge per residue: positive (blue) or negative (red); I) Das-Pappu phase [29] (plot shown in M): globular (1, green), Janus/boundary (2, yellow), strong polyelectrolyte (3, purple), strong polyanion (4, red), strong polycation (5, blue); J) dSNP enrichment (blue) or depletion (red) based on analysis using human SNPs [2] (plot shown in M); K) position on the Uversky-Gillepse-Fink boundary plot [30] (plot shown in M); and L) predicted fraction of disordered residues, according to PV2 via D2P2. M) From left to right: legends corresponding to the coloring schemes for the tracks in I [29], J (used with permission from [2], Copyright 2022 PNAS), and K [30]. The example protein shown in these tracks is insulin (UniProt ID: P01308).

After the initial blobulation, the user can interactively tune some of the parameters. For example, while our previous studies [1, 2], the example applications in this paper, and the webtool default use the Kyte-Doolittle scale [31], the user may select the Eisenberg-Weiss scale [32] or the Moon-Fleming scale [33]. Additionally, the user can change the hydropathy scale used for digitization (Fig. 2A) or interactively modify the *H*^∗^ threshold and *L*_min_ cutoff (here we use *H*^∗^= 0.4 and *L*_min_= 4, respectively) to segment the protein into modules with varying hydropathy and length properties. This can be done by adjusting the respective sliders or manually entering a value (Fig. 2B and C). A user may also introduce a mutation by selecting one of the black triangles or manually entering the position and alternate amino acid in the “mutate residue” field and selecting the checkbox (Fig. 2D). All changes update dynamically. Users can download both an image of the output as a PDF and the raw data (in CSV format) used to generate the webtool output (Fig. 2E).

The smoothed hydropathy of each residue *H_i_* (as defined in section 2) is shown in the first “Results” track (Fig. 2F) along with the *H*^∗^ threshold (blue line). While a graph displaying the hydropathy of individual residues is common to many tools, blobulation clusters adjacent residues by their hydrophobicity, identifying blobs. The subsequent tracks display blobs colored by various biochemical properties, and more a detailed description is provided in the SI.

### 3.2 Python Scripting and Command Line Interface

For use in high-throughput applications, like those presented in Ref. 2, we also provide a stand-alone Python package for blobulation. This package is also the backend for the *blobulator* webtool, and is called when a protein is submitted via that interface. The package can be used either in Python scripts or directly via a command line interface (CLI), which is pip-installable:

> pip install blobulator

The input to the CLI can be either amino acid or DNA sequences in plain text or FASTA format , the latter of which is specified with -Fasta or -DNA flags, respectively. For a plain text sequence:

> python3 -m blobulator --sequence AFRPGAGQPPRRKECTPEVEEGV --oname

> *’*→ ./my_blobulation.csv

For a FASTA file:

> python3 -m blobulator --fasta ./relative/path/to/my_sequences.fasta --oname

> *’*→ ./relative/path/to/outputs/

The outputs of the CLI are CSV files (one for each input sequence, in the above example, this is named my blobulation.csv) identical to the downloadable blobulation output of the webtool (see 3.1). Additional example scripts are provided on the *blobulator* GitHub.

### 3.3 VMD Plugin

While the webtool allows users to view the blobulation of the sequence and various averaged physicochemical properties, the plugin tool for Visual Molecular Dynamics (VMD) [19] allows users to incorporate blob information into the visualization and analysis of protein structure and dynamics. Users can assign blobs with the same parameters and algorithm in the webtool, while taking advantage of the existing VMD tools for scriptable analyses and creation of high-quality images and movies. Additionally, the blobulation algorithm was reimplemented in TCL, the native VMD scripting language, for maximum compatibility. The TCL script extracts the sequence information from the provided structure and then detects blobs based on user-provided parameters. The user, user2, and user3 fields associated with each atom store the type of blob (h, p, s), the blob ID, or the group ID, respectively. This information can then be used in atom selections or user-field coloring schemes provided by the VMD software; the molecular images in this manuscript were created by creating an atom selection for each h-blob, representing it with “quick-surf”, and coloring each h-blob a slightly different shade of blue. Examples of images created using this plugin tool can be found throughout applications in this paper (Figs. 4 and 5). For proteins with many blobs, this approach can require many separate graphical representations; we include an automated approach to create the representations in a batch process.

Since users of VMD typically create representations through GUIs, we also provide a GUI through which the user can provide and adjust parameters, as well as call the scripts that blobulate the sequence and create the representations. This GUI is similar to the control panel found on the *blobulator* webtool’s output page, with some additional features. There is an option to blobulate only part of a protein using the Selection field, which may be useful for large proteins or to “zoom in” on a given set of residues (Fig. 3B). Adjacent to the threshold sliders, there are buttons to restore each field to its default value: 4 and the equivalent of 0.4 on the selected scale for the length and hydropathy thresholds, respectively (Fig. 3D and E). Additionally, in the viewer window, users are able to adaptively tune parameters and view how this changes the blobs found on a protein’s structure. Finally, using a dropdown menu (Fig. 3F), users can color blobs by their type (all h-blobs are colored the same shade of blue), or by their ID (all h-blobs are colored distinct shades of blue). Scripts and installation instructions for the VMD plugin can be found in the *blobulator* GitHub repository: https://github.com/BranniganLab/blobulator.

**Figure 3.**
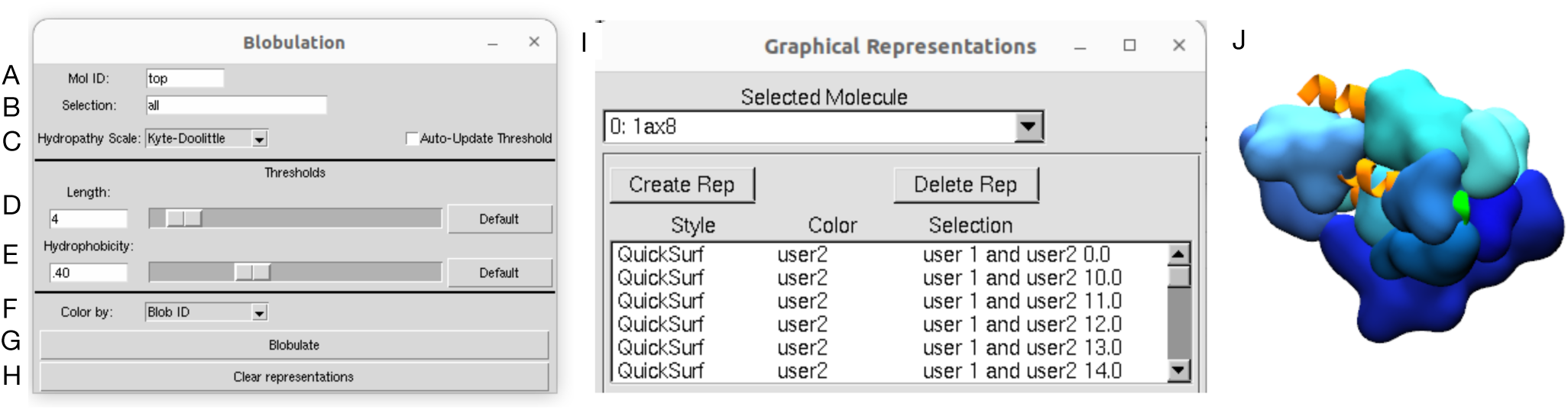
Screenshot of the VMD blobulator plugin. The user sets the Mol ID of the protein (A), provides an atomselection for the display (B), selects a hydropathy scale and may select the auto update option to snap the threshold to a scale-dependent default (C), sets the minimum blob length *L*_min_ (D) and hydropathy cutoff *H*^∗^ (E), selects how blobs are colored (F). The “Blobulate” button initiates blobulation and creates graphical representations within VMD (G). All representations are deleted by clicking the “Clear representations” button (H). The “Graphical Representations” control panel contains the created graphical representations, which can be further modified (I), and are eventually displayed in the example viewer window (J) showing Leptin (PDB: 1ax8).

## 4 EXAMPLE APPLICATION: LYSOZYME

In this section, we present the blobulation of lysozyme, and demonstrate how blobulation can provide a hydrophobicity-based framework for identifying protein modularity using only the sequence. We have chosen an example protein with previously established features, such as binding sites and disease-associated mutations, and illustrate the context that blobulation provides to these known features and how future research might utilize the *blobulator* toolkit for proteins with less established features. Additional example applications are presented in the SI.

In an aqueous environment, most globular proteins have a highly hydrophobic core surrounded by a solvent-accessible surface. One such protein is lysozyme, which cleaves the sugar and peptide components of peptidoglycan. To provide an example of how one might detect hydrophobic blobs that correspond to the hydrophobic core of a globular protein, we blobulated bacteriophage T4 lysozyme and varied both the hydropathy threshold (*H*^∗^) and minimum length (*L*_min_) (Fig. 4). Figure 4 B shows a structure of lysozyme colored by blob type, and we note that h-blobs do not align with secondary structure elements. Lysozyme has two catalytic residues near the N-terminus and several substrate contact sites along the sequence (Fig. 4A). Higher hydropathy and length thresholds eliminate the h-blobs that are detected at the surface of the protein when using more relaxed settings (Fig. 4C). Blobulation using the most stringent settings shown here (*H*^∗^= 0.5, *L*_min_= 8) reveals two h-blobs at the center of the protein, away from the solvent-accessible surface. By gradually increasing parameter thresholds, we can isolate the components of a globular protein that correspond to the core, as well as the shorter and less hydrophobic blobs that interact at the surface of the protein. This is consistent with previous findings that h-blobs tend to be buried in structured proteins [2].

To investigate whether the substrate binding site is composed of h-blobs, we identified blobs containing contact residues within 7 Å of peptidoglycan (shown in Fig. 4). When using a shorter *L*_min_ (*L*_min_= 4), h-blobs are found contacting peptidoglycan across its entire length. However, when using a longer *L*_min_ (*L*_min_= 8), the only detected h-blobs in the substrate binding site are those in contact with the peptidoglycan sugar component. Stabilization of this sugar ring is vital for the ability of the enzyme to cleave the peptidoglycan [36], and we find this ring wedged between two long hydrophobic blobs (Fig. 4C), providing an example of long hydrophobic blobs with interactions critical for function.

**Figure 4.**
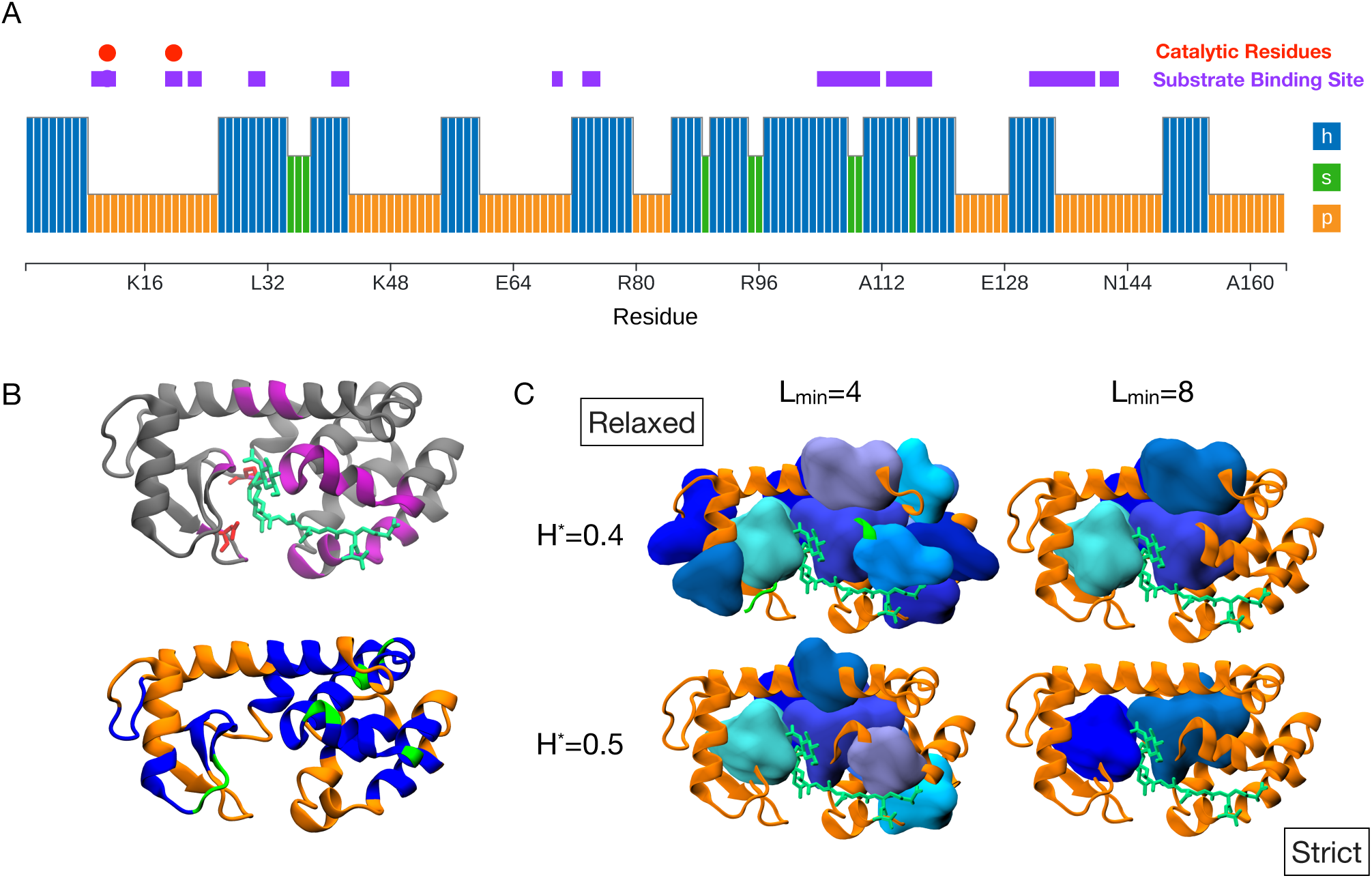
Blobulation of lysozyme. A) Blobs colored according to blob-type, as outputted from the *blobulator* webtool, and produced using default settings (*H*^∗^= 0.4, *L*_min_= 4). Annotations indicate catalytic residues (red) and the substrate binding site (purple). B) Molecular image of lysozyme (PDB:148L) with peptidoglycan (green) colored by the substrate binding site (left, residues found within 7 Å of peptidoglycan) or by blob type (right, h-blob: blue, p-blob: orange, s-blob: green). C) Blobulation under increasingly stringent settings (*H*^∗^= 0.4 and 0.5, *L*_min_= 4 and 8). H-blobs are shown as surfaces. Molecular images were generated in VMD [34, 35] using the VMD plugin introduced in 3.3.

Blobulating lysozyme using “relaxed” settings (*H*^∗^= 0.4 and *L*_min_= 4) reveals two examples of “blob groups”: sets of h-blobs separated only by s-blobs (as defined in section 2, shown in gray in Fig. 5A). Both blob groups are found near the center of lysozyme, surrounded by ungrouped h-blobs and p-blobs. Additionally, each group contains an h-blob that remains detected under increasingly stringent settings and is also found at the core of lysozyme (Fig. 5B, shown also in the molecular image in 4). The groups detected here are akin to tertiary elements formed from a network of secondary interacting structure elements, such as alpha-helical bundles. This is an example of hierarchical blob clustering, where hydrophobic blobs detected at restrictive parameters are found in the protein core surrounded by less hydrophobic blobs within the same group, which are in turn surrounded by individual h-blobs.

**Figure 5.**
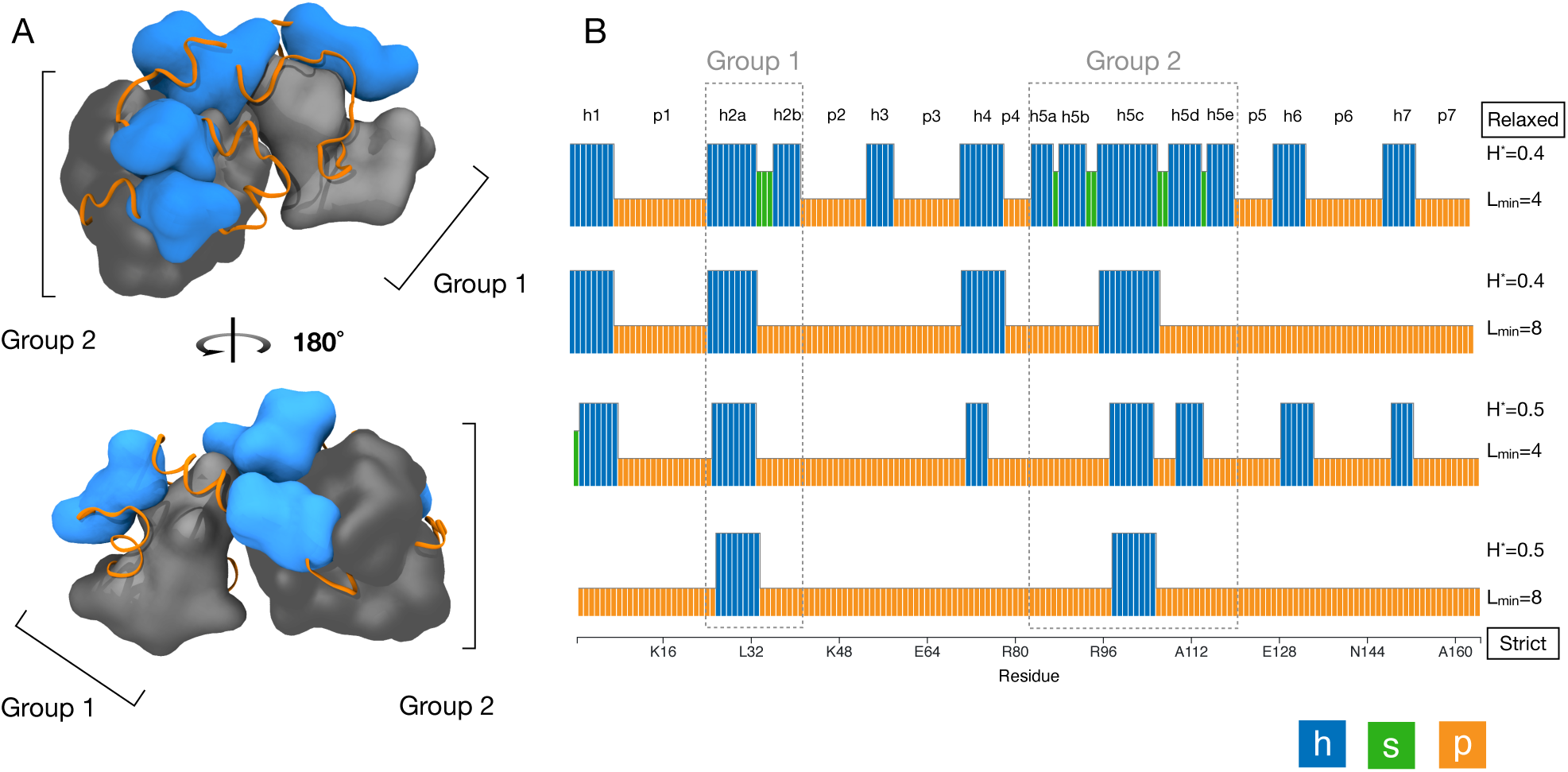
Blob groups in T4 lysozyme. A) Structural view (PDB:2LZM) blobulated using default settings (*H*^∗^= 0.4, *L*_min_= 4). Groups are gray, ungrouped h-blobs are blue, and p-blobs are orange ribbons. B) Blobulation of T4 lysozyme under increasingly stringent settings. Annotations indicate blob identifiers and blob groups. Molecular images were generated in VMD [34, 35].

To provide an example of how blobulation links known mutations with an effect on their local hydrophobic context, we blobulated the lysozyme mutants S117V and T157I. Both mutations affect the temperature on stability of the lysozyme by altering intraprotein hydrophobic interactions. S117 makes the protein more stable at higher temperatures by altering hydrophobic residue interactions in the substrate binding cleft [37], while T157I makes the protein less stable at higher temperatures and disrupts hydrogen bonding at the periphery of the protein [38]. We find that both mutations change the blob topology (Fig. 6): S117V merges two h-blobs, and T157I creates a new h-blob four residues in length. We have previously found that mutations that split, dissolve, or merge h-blobs are enriched for deleterious mutations [2], and this result is consistent with that finding. Additionally, the h-blob introduced by the T157I mutation is classified as a Janus region on the Das-Pappu phase diagram [29]. Janus proteins often have degenerate conformations and switch between ordered and disordered depending on their environment [29], which may cause a change in this blob at higher temperatures and lead to less overall stability for the protein. Finally, single-residue mutations frequently do not cause detectable structural differences in predictions; for instance, AlphaFold predicts minimal structural differences between these mutants and the wild type (RMSD *<* 0.1 Å). In contrast, the sequence-based topology from blobulation is sensitive to these single-residue mutations.

**Figure 6.**
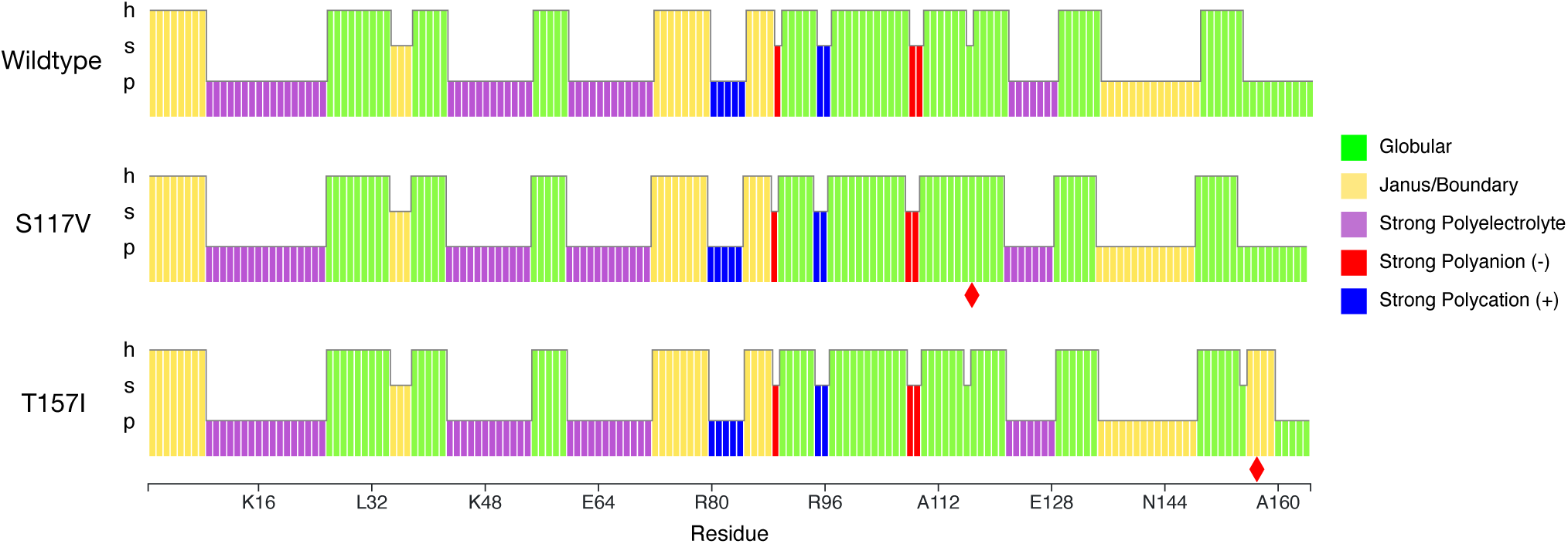
Globular tendency of T4 lysozyme blobs. Blobulation using default parameters (*H*^∗^= 0.4, *L*_min_= 4). Blobs for each sequence are colored by Das-Pappu phase [29], as in Fig. 2I. Red diamonds indicate mutated residues. S117V, which increases thermostability, joins two h-blobs into one. T157I, which decreases thermostability, creates a new h-blob (as well as an s-blob).

## 5 CONCLUSION

Here we presented the *blobulator* toolkit for characterizing and visualizing patterns of contiguous hydrophobicity in proteins. Blobulation reflects higher-level sequence organization, much like secondary structure elements, but applies even when structural data are unavailable. The runtime of the CLI is comparable to modern secondary structure predictors and scales about four times more efficiently with sequence length. The webtool and VMD viewer provide a graphical and interactive means to explore the hydrophobicity in a single protein in depth, including blob-level properties like net charge and disorder, as well as the impact of amino acid substitutions.

As illustrated in the example applications and in Ref 2, sequence partitioning by hydrophobic blobs is compatible with known functional segments and tertiary interactions. Tertiary interactions are increasingly incorporated into bioinformatics analyses [13, 39–46] typically through machine-learning approaches. Since hydrophobic blobs are critical determinants of tertiary interactions, the *blobulator* CLI suggests a deterministic, lightweight, and biophysically-informed route to bioinformatics analyses. For example, this toolkit opens the door to studies that require high statistical power across multiple sequences, such as the long-term evolution of protein hydrophobicity or the impact of disrupting hydrophobic blobs for genetic disease.

## 6 DATA AND SOFTWARE AVAILABILITY STATEMENT

The backend for the *blobulator* toolkit can be found on GitHub at https://github.com/BranniganLab. Additionally, the webtool is available as a graphical user interface at https://www.blobulator.branniganlab.org/.

## 8 AUTHOR CONTRIBUTIONS

CP, ES, RL, RWL, MEBH, and GB conceived of the project. CP, ES, RL, RWL, KB, LR, and TJ contributed to toolkit development. CP, ES, RL, and GB wrote the paper. TJ MH, and GB supervised.

## 9 ACKNOWLEDGEMENTS

The authors acknowledge the Office of Advanced Research Computing (OARC) at Rutgers, The State University of New Jersey, for providing access to the Amarel cluster and associated research computing resources. TJ is supported by NIH 1K08GM139031, MEBH is supported by NIH 1R35GM134957, and LR, ESM, CP, and GB are supported by NSF DGE 2152059. We are also grateful to Dr. Jérôme Hénin, Mr. Jesse Sandberg, and Mr. Jahmal Ennis for testing and feedback.

