## Supporting Information for "The blobulator: a toolkit for identification and visual exploration of hydrophobic modularity in protein sequences"

### Supplementary Methods

#### *Toolkit Overview*

The *Blobulator* toolkit is comprised of a command line interface (CLI), a webtool, and a VMD plugin. The accepted inputs for each tool at the time of writing are shown in Figure S1. We note that an option to provide structures as an input for the webtool is in development.

#### *Installation and Usage Instructions for the Blobulator Toolkit*

**Blobulator Command Line Interface**
**Software requirements:** Python 3.9+

**Quick Install:** (Optional) Create a conda environment:

```
conda create --name blobulator_env python=3.9
conda activate blobulator_env
```

**Download the repository:**

```
git clone https://github.com/BranniganLab/blobulator
```

**Install with pip:**

```
pip install git+https://github.com/BranniganLab/blobulator
```

**Scripting Example (Python):**

```python
import blobulator

# A very simple oligopeptide and standard settings
sequence = "RRRRRRRRRIIIIIIIII"
cutoff = 0.4
min_blob = 4
hscale = "kyte_doolittle"

# Do the blobulation
blobDF = blobulator.compute(sequence, cutoff, min_blob, hscale)

# Cleanup the dataframe (make it more human-readable)
blobDF = blobulator.clean_df(blobDF)

# Save it as a csv for later use
oname = "hello_blob.csv"
blobDF.to_csv(oname, index=False)
```

Additional sample scripts can be found in the repository examples directory.

**Using the command-line utility:**

*Basic usage:*

```
python3 -m blobulator --sequence AFRPGAGQPPRRKECTPEVEEGV --oname
 ↪  ./my_blobulation.csv
```

This will blobulate the sequence ”AFRPGAGQPPRRKECTPEVEEGV” and write the result to my_blobulation.csv.

*Advanced Usage (FASTA files):*

Place a fasta file with one or more sequences in any directory (they must all be DNA or protein sequences), then open a terminal in the blobulator directory and run:

```
python3 -m blobulator --fasta ./relative/path/to/my_sequences.fasta --oname
 ↪  ./relative/path/to/outputs/
```

This will blobulate all sequences in my_sequences.fasta (assuming they are protein sequences) and output the results to the outputs folder prefixed by their sequence ID.

**Blobulator VMD Plugin**
**Software requirements:** VMD

**Installation guide:**
To obtain this plugin, download the following files from the VMD_scripts folder into a single directory:

- blobulation.tcl
- Blob_GUI.tcl
- normalized_hydropathyscales.tcl

**Quickstart:**

1. Load a protein into VMD.
2. Access the Tk console via the Extensions dropdown menu ‘Extensions → Tk Console‘.
3. In the Tk console, change directory to the directory where you downloaded the above scripts: (cd /path/to/blobulator/scripts).
4. Source the plugin (source Blob_GUI.tcl‘).
5. Click the blobulate button to generate the corresponding graphical representation in VMD.

**How to access blob representations in VMD:**

- The blobulation algorithm will apply all blob types to the VMD user and user2 values.
- The ’user’ value will store the type of blob: user 1 → h-blobs, user 2 → s-blobs, and user 3 → p-blobs.
- The ’user2’ value will store the blob group: user2 1 → h-blob group 1, user2 2 → s-blob group 1, user2 3 → h-blob group 2, etc.
- When coloring by Blob ID, h-blobs will have different colors depending on the user2 value.

**Blobulator Webtool**

**Quickstart:**

1. The webtool can be found at `http://www.blobulator.branniganlab.org/`
2. From the New Query page, select a protein to blobulate, either via:
   (a) ID Entry (for Uniprot or Ensembl IDs)
   (b) Manual Entry (for protein sequences)

3. Click the ’Blobulate’ button
4. On the outputted Result Page, toggle blobulation settings, and view tracks showing blobs colored by various biochemical properties.
5. Download the underlying data, or print the screen to save a version of the blobulation output or tracks, respectively.

#### ***Pseudocode Blobulation Algorithm***

While the blobulation algorithm was described in section 2, we have included a pseudocode representation here.

##### Digitization

In step 1 of the algorithm, the hydrophobicity of each residue is digitized and smoothed. The required inputs for this step are a hydrophobicity scale (hydroscale), a hydrophobicity threshold ($H_{star}$), a length minimum ($L_{min}$), and a protein sequence (sequence).

```
sequence_length = length(sequence)
residue_hydrophobicities = empty list of length sequence_length
digitized_residues = empty list of length sequence_length
for i = 1 to sequence_length do
    residue_hydrophobicities[i] = hydroscale[sequence[i]]
end for
for i = 1 to sequence_length do
    if i == 1 then
        smoothed_hydrophobicity = (residue_hydrophobicities[i] + residue_hydrophobicities[i + 1]) / 2
    else if i == sequence_length then
        smoothed_hydrophobicity = (residue_hydrophobicities[i] + residue_hydrophobicities[i - 1]) / 2
    else:
        smoothed_hydrophobicity = (residue_hydrophobicities[i – 1] + residue_hydrophobicities[i] +
            residue_hydrophobicities[i + 1]) / 3
    end if
    if smoothed_hydrophobicity > H_star then
        digitized_residues[i] = “hydrophobic”
    else
        digitized_residues[i] = “non-hydrophobic”
    end if
end for
return digitized_residues
```

##### Clustering

After all residues are digitized, they are clustered into blobs. The digitized_residues, sequence_length, and length minimum ($L_{min}$) variables from step 1 are used as the input for step 2, in which residues are clustered into blobs. The blobulated sequence is returned.

```
hydrophobic_subseq = empty string
non-hydrophobic_subseq = empty string
blobulated_seq = empty string
for i = 1 to sequence_length do
    if digitized_residues[i] == ”non-hydrophobic” then
        if length(hydrophobic_subseq) ≥ L_min then
            blobulated_seq += hydrophobic_subseq
        else if length(hydrophobic_subseq) != 0 then
            non-hydrophobic_subseq += “p” * length(hydrophobic_subseq)
            non-hydrophobic_subseq += ”p”
        end if
        hydrophobic_subseq = empty string
    end if
    if digitized_residues[i] == “hydrophobic” then
```

```
        if length(non-hydrophobic_subseq) ≥ L_min then
            blobulated_seq += non-hydrophobic_subseq
        else if length(non-hydrophobic_subseq) != 0 then
            blobulated_seq += "s" * length(non-hydrophobic_subseq)
        end if
        hydrophobic_subseq += "h"
        non-hydrophobic_subseq = empty string
    end if
    if i == sequence_length then
        if length(hydrophobic_subseq) ≥ L_min then
            blobulated_seq += hydrophobic_subseq
        else if length(non-hydrophobic_subseq) ≥ L_min then
            blobulated_seq += non-hydrophobic_subseq
        else if length(non-hydrophobic_subseq) != 0 then
            blobulated_seq += "s" * length(non-hydrophobic_subseq)
        end if
    end if
end for
return blobulated_seq
```

##### *Choosing $H^*$ and $L_{min}$*

While we note that there are no “correct” parameters for blobulation, we have included some example settings as a starting point for a few research applications. These are provided in table S1.

##### *Webtool Track Descriptions*

The track below “Smoothed hydropathy per residue” shows the output of the clustering step (Fig. 2G), where blobs are colored by their blob type. Blobulation of insulin using the settings $H^*$= 0.4 and $L_{\text{min}}$= 4 identifies 8 h-blobs, 5 p-blobs, and 3 s-blobs. If a UniProt ID is provided, known disease-associated mutations are obtained from UniProt and displayed on every track as black triangles (Fig. 2). Hovering over these with the cursor will display the effect of the mutation on the amino acid, its reference SNP cluster ID (rsid), and link to the Single Nucleotide Polymorphism Database (dbSNP).

The webtool includes a series of tracks in which blobs are colored by additional information relating to charge composition, hydrophobicity, length, or some combination of the three. As shown in Figure 2H-L, these include tracks displaying blob charge, globular tendency, the predicted propensity of each blob to deleterious mutations, and two tracks showing order and disorder predictions for each blob, respectively.

The first two of these tracks are colored by mean net charge per residue and globular tendency, both calculated based solely on charged residues in the blob. (Fig. 2H and 2I). In the mean net charge per residue track (Fig. 2H), blobs are colored by the average charge of all the residues within each blob. The use of this track is demonstrated in the example in Membrane Proteins: Pentameric Ligand-Gated Ion Channels (pLGICs) below. In the globular tendency track (Fig. 2I), blobs are colored based on the region they occupy in the Das-Pappu diagram (reproduced in Fig. 2M). Traditionally, the Das-Pappu phase is the globular tendency based on the charge content of a whole protein sequence [1], but here we have calculated the Das-Pappu phase of each blob. The use of this track is demonstrated in section 4.

The predicted mutation sensitivity track is colored based on an enrichment calculation that includes both blob hydrophobicity and length (Fig. 2J). This track colors blobs based on the enrichment of disease variants in blobs with the same properties found in our previous study [2]. Enrichments were calculated from a dataset containing about 70,000 human disease-associated and non-disease-associated missense mutations (shown in Fig. 2M, used with permission from [2], Copyright 2022 PNAS).

Finally, we provide two tracks that report the estimated disorder for the subsequence of amino acids contained in each blob (Fig. 2K and 2L). We note that blob-type is expected to be, at most, weakly correlated with disorder and these tracks do not incorporate the blob type in the disorder prediction. Instead, these tracks provide an approach for intermediate-level aggregation of residue-level disorder, yielding higher resolution than a whole-sequence estimate.

The Uversky-Gillespie-Fink equation for the boundary between disordered and ordered proteins considers only aggregate properties of a sequence (the average hydropathy $\langle H\rangle$ and the net charge $\langle R\rangle$), and is given by [3]

$$\langle H\rangle = a\langle R\rangle + b, \tag{1}$$

where $a = 0.359$ and $b = 0.413$. Eq. 1 neglects the order of residues within the sequence, making it most appropriate for short sequences. In the track shown in Fig. 2K, we use this metric to estimate the disorder for each blob: the signed distance from this boundary for blob $i$ with average hydropathy $\langle H\rangle_i$ and net charge $\langle R\rangle_i$ is given by

$$d = \begin{cases} \sqrt{\frac{(\langle H\rangle_i - a\langle R\rangle_i - b)^2}{1+a^2}}, & \text{if } \langle H\rangle_i - a\langle R\rangle_i - b > 0 \\ -\sqrt{\frac{(\langle H\rangle_i - a\langle R\rangle_i - b)^2}{1+a^2}}, & \text{otherwise.} \end{cases}$$

where $d > 0$ and $d < 0$ correspond to more ordered and more disordered blobs, respectively. Each blob is colored according to the associated value of $d$, with the 2-dimensional color scale shown in Figure 2M. This approach preserves longer-range patterns of order and disorder, while applying a deterministic, pattern-independent estimate of disorder to each chunk.

While the track shown in Fig. 2K aggregates the residual information in the blob before estimating disorder, the next track (Fig. 2L) estimates disorder per residue and then aggregates it across the blob. The predicted disorder by residue for a given protein sequence, measured by PV2 [4], is accessed from Uniprot (only available if 'ID Entry' is used). Each blob is then colored by the fraction of contained residues predicted to be disordered.

### Supplementary Results

#### *Membrane Proteins: Pentameric Ligand-Gated Ion Channels (pLGICs)*

Unlike soluble globular proteins, which enclose their hydrophobic residues inside a hydrophilic shell, integral membrane proteins use exposed hydrophobic regions as anchors in the cell membrane. The pentameric ligand-gated ion channel (pLGIC) protein family is responsible for neurotransmitter reception in the nervous system of bilateria, but representatives are also found in many other eukaryotes and some prokaryotes [5]. Despite relatively low sequence identity and diverse environmental and ligand requirements, pLGICs are highly structurally conserved [6–10]. The N-terminal extracellular domain consists of a beta-sandwich, which is responsible for ligand sensing and much of the substrate selectivity (Fig. S2A-B). The transmembrane domain consists of four transmembrane helices per subunit (Fig S2D)[5]. When the ligand is bound in the ECD, a signal is transmitted to the transmembrane helical bundle (Fig. S2D), which changes conformation to permit ions to pass through the channel formed by the five pore-lining helices. In this example, we will compare the blobulation of two structural models of the pLGIC family: first, the *Caenorhabditis elegans* glutamate-gated chloride channel (GluCl); and second, the proton-gated *Gloeobacter* ligand-gated cation channel (GLIC).

Starting from the N-terminus, both GluCl and GLIC contain disordered N-terminal extensions which include signal sequences (Fig. S3). GluCl has an additional hydrophobic blob in the N-terminal extension. P-blobs dominate the extracellular beta-sandwich in both cases. The two additional hydrophobic blobs in GluCl correspond to $\beta 1$ and $\beta 5$, which contain two of the four residues that contact glutamate during binding. GLIC, in contrast, is proton-gated via titratable residues mostly in the ECD [7] and lacks these extracellular h-blobs. The transmembrane helices are clearly visible using $H^* = 0.33$ and $L_{\text{min}} \approx 19$. This length corresponds to the typical thickness of a plasma membrane[11]. The extended p-blob seen in the blobulation of GluCl (Fig. S2) corresponds to a disordered intracellular domain common in eukaryotic pLGICs [7].

As ion channels, the electrostatic potential of a pLGIC pore plays a central role in both conductivity and selectivity. The blobs along the ion conduction pathway (in the ECD and the M2 helix) of GluCl are primarily positive or neutral. GLIC, in contrast, has a net negative charge in these regions (Fig. S3). This is consistent with the anion conductivity of GluCl and the cation conductivity of GLIC. The charges in the M2 blob are on the intracellular side, close to the M1-M2 loop. GluCl has a single positive charge (R306), while GLIC has several negative charges in this region. GLIC has an additional positive charge near the M2-M3 loop due to K290, which can be seen by using the blobulator's zoom feature. Finally, the M4 helix of each protein has a net positive charge, consistent with experimental observations that M4 binds anionic lipids [12, 13]. These results show the blobulator's potential for extracting structural information from membrane proteins either individually or within a larger evolutionary context.

#### *IDP: $\alpha$-synuclein*

Intrinsically disordered proteins (IDPs) fulfill critical roles despite lacking stable tertiary structure (as reviewed in Ref. 14). One such IDP is $\alpha$-synuclein. Although its function is not fully understood, it forms aggregates implicated in Parkinson's disease [15, 16]. It contains a helix-turn-helix motif at its N-terminal and a random coil near its C-terminal. Although residues throughout the helical region of this protein interact with other $\alpha$-synucleins, residues 71 to 82 are the necessary aggregating motif [17] (Fig. S5B). To identify whether h-blobs overlap with this known motif, we blobulated $\alpha$-synuclein using the settings $H^*= 0.4$ and $L_{\text{min}}= 4$ and found that the h-blob group in $\alpha$-synuclein coincides with this aggregating motif (Fig. S5A). This result demonstrates that the blobulation of IDPs, where secondary structure elements are often transient, can reveal hydrophobic clusters that in some cases coincide with tertiary interactions.

One application for blobulation is in quantifying the effects of mutated residues on their local sequence context. The helix-turn-helix motif of $\alpha$-synuclein (Fig. S5B) also interacts with cell membranes [18], but the mechanism driving this interaction is unknown [18]. Additionally, while this motif contains the aggregating region (pink), the two longest h-blobs detected under these settings overlap the majority of it, extending past the edge of the alpha helix and into the random coil. Here, we consider the known disease-causing mutations A30P and A53T. When expressed in yeast, A53T and wildtype $\alpha$-synuclein initially bind to the membrane before forming cytoplasmic aggregates, whereas A30P $\alpha$-synuclein is dispersed throughout the cytoplasm [19]. The A30P mutant has a shorter and less ordered third h-blob than A53T and wildtype $\alpha$-synuclein (Fig. S4C). Because the order prediction considers residue hydrophobicity, and proline is less hydrophobic than alanine, a decrease in order is expected because predicted order increases with hydrophobicity on the Uversky-Gillepse-Fink boundary plot [3]. This is also consistent with the finding that while the helical domain of the mutant is partially disrupted, its N-terminus is more dynamic than the wildtype [20]. In contrast to the wildtype, no blobs are shortened in the A53T mutant protein, and the order of the blob containing the mutated residue is only minimally altered.

### Supplementary Tables

| **Research Application** | $H^*$ | $L_{min}$ |
|---|---|---|
| IDPs/General Use | 0.40 | 4 |
| Hydrophobic Core | 0.50 | 8 |
| Transmembrane Domains | 0.33 | 19 |

**Table S1.** Example starting settings for $H^*$ and $L_{min}$ by research application. We recommend first tuning $H^*$, followed by $L_{min}$.

### Supplementary Figures

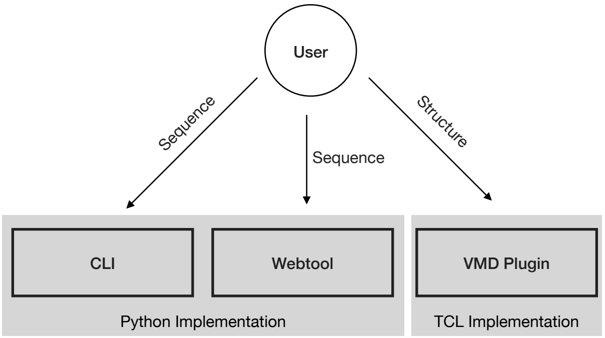


**Figure S1.** **Diagram showing tools in the *Blobulator* toolkit.** Accepted inputs are shown as labeled lines to each tool. The language that the backend implementation of the blobulation algorithm for each respective tool is shown as a gray box.

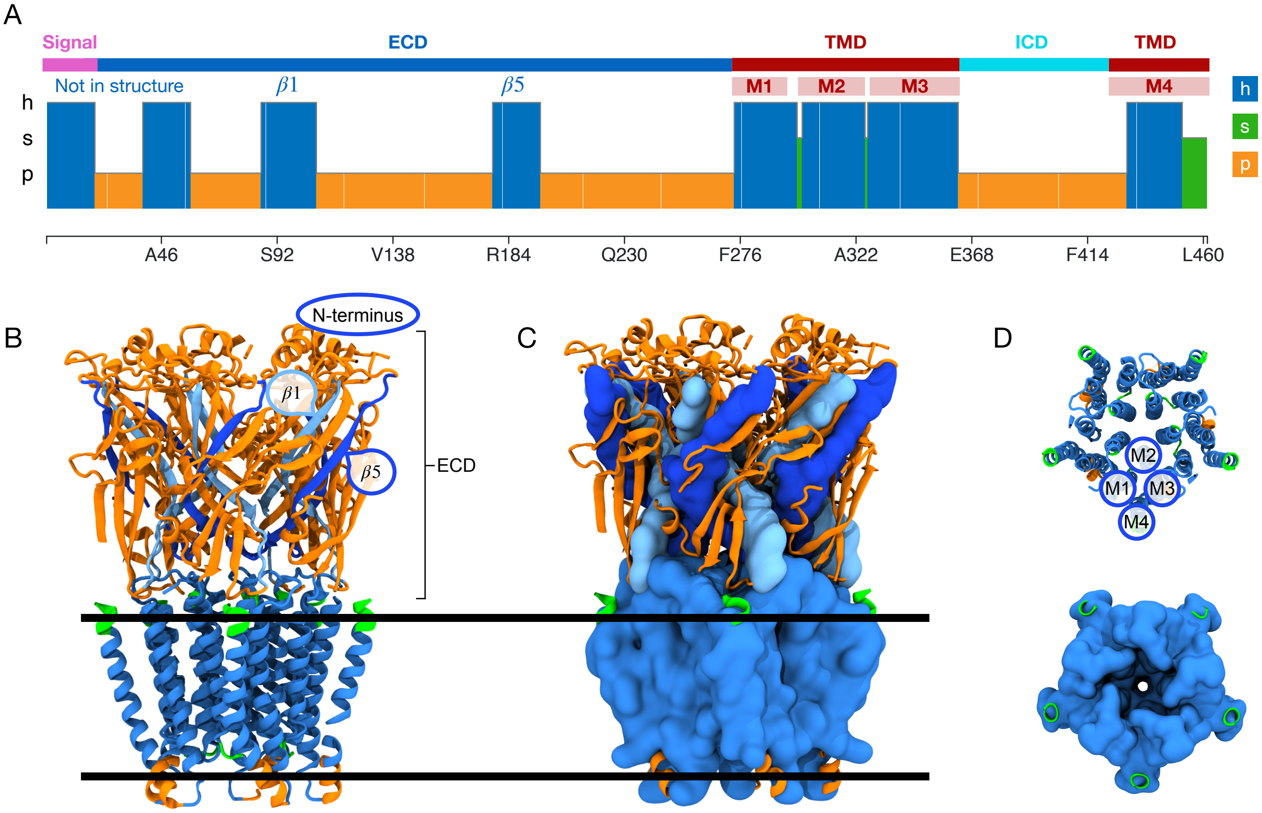


**Figure S2. Blobulation of GluCl, a pentameric ligand-gated ion channel.** A) Blobulation of GluCl (UniProt ID: G5EBR3) using settings to detect transmembrane regions ($H^* = 0.33$, $L_{\text{min}} = 19$). B) A cartoon representation of GluCl (PDB: 3RHW) colored according to blob type and ID: $\beta 1$ blob (light blue), $\beta 5$ blob (dark blue), transmembrane blobs (intermediate blue), terminal s-blobs (green), p-blobs (orange). The first two blobs in the sequence are absent from the structure. The black lines represent the lipid membrane. C) GluCl showing h-blobs in surface view to better show blob-blob contacts. Colored as in B. D) Extracellular views of the TMDs of the proteins shown in B (upper) and C (lower). Molecular images were generated in VMD [21, 22].

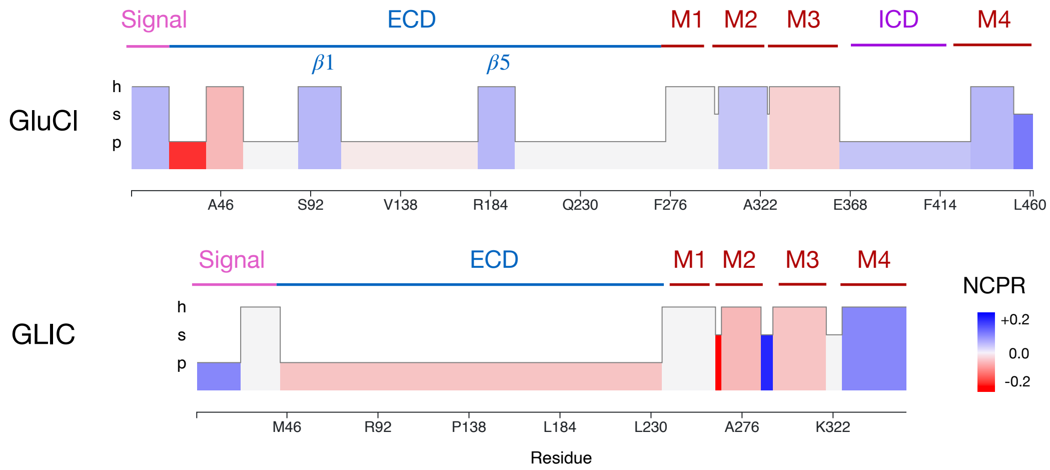


**Figure S3.** **Comparative blobulation of GluCl (as in Fig. S2), an anion-conducting pLGIC, and GLIC, a cation-conducting pLGIC.** Net Charge Per Residue (NCPR) tracks for GluCl (UniProt: G5EBR3) and GLIC (UniProt: Q7NDN8) blobulated using settings to detect transmembrane regions ($H^* = 0.33$, $L_{\text{min}} = 19$). Tracks are aligned by the beginning of the h-blob containing the M2 (pore-lining) helix. Annotations indicate important sequence features available on UniProt and indicated in Fig. S2. Signal sequences are indicated in pink, ECD regions are indicated in blue, and transmembrane helices (M1 to M4) are indicated in red.

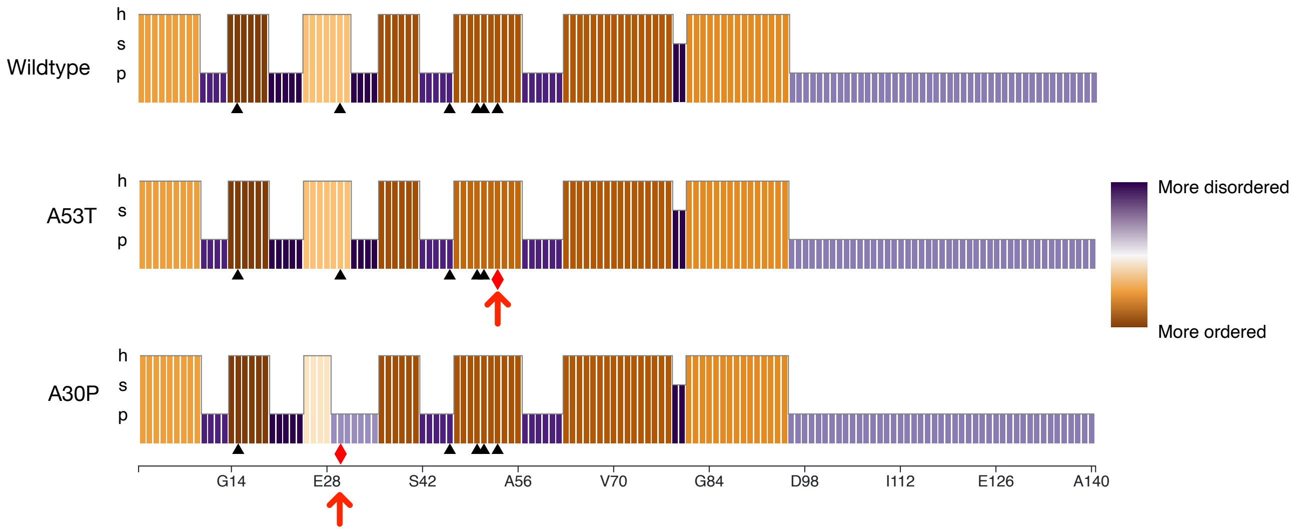


**Figure S4. Order predictions for blobs of $\alpha$-synuclein.** Wildtype, A53T, and A30P mutants colored by blob disorder as calculated using each blob's signed distance from the order/disorder boundary of the Uversky-Gillepse-Fink boundary plot [3]. Mutations are indicated: A53T and A30P (red diamonds and arrows), and other known disease-associated mutations (black triangles). Blobulation uses default settings ($H^{*}$= 0.4, $L_{\text{min}}$= 4).

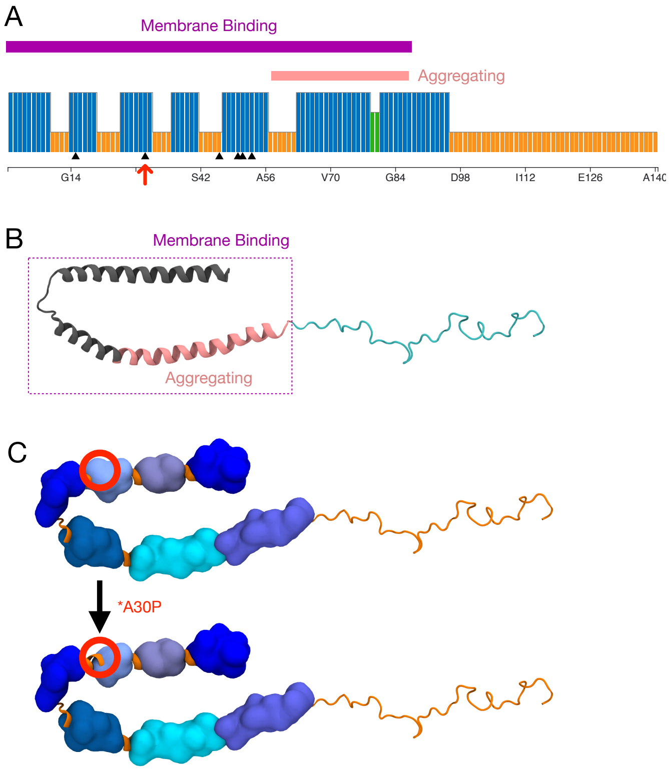


**Figure S5. Blobulation of $\alpha$-synuclein, an intrinsically disordered protein.** A) Webtool "blob type" track of $\alpha$-synuclein (UniProt ID: P37840) blobulated using default settings ($H^*$= 0.4, $L_{min}$= 4). Annotations indicate the membrane-interacting region (purple) and the protein-interacting region (pink). B) $\alpha$-synuclein structure (PDB: 1XQ8), labeled by the membrane-interacting region in cartoon. Aggregating in pink, non-aggregating in purple. C) Changes to blobs (blue surfaces, change indicated by red circles) caused by the A30P mutant compared to the wildtype. Molecular images were generated in VMD [21, 22].

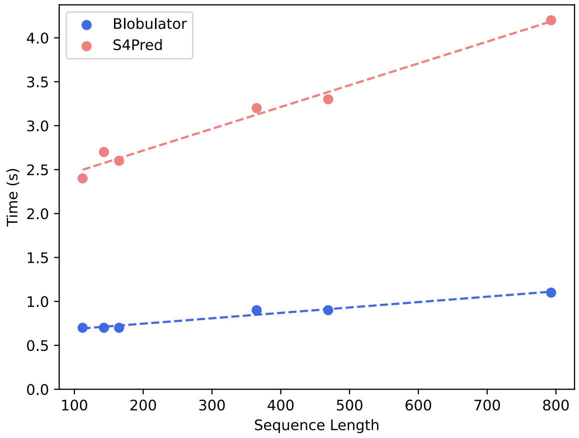


**Figure S6. Runtime of *blobulator* CLI (blue) and secondary structure prediction tool S4Pred[23] (red) as a function of protein sequence length.** Six sequences were selected for the benchmarks to cover a range of protein lengths. Benchmarks were run on a single core of a workstation (CPU: Xeon W1390). Each sequence was processed ten times and the average is reported. The dashed lines represent linear least squares fits for the *blobulator* (slope: 0.00061, y-intercept: 0.62) and S4Pred (slope: 0.0025, y-intercept: 2.2).

[20] Luisel R. Lemkau, Gemma Comellas, Kathryn D. Kloepper, Wendy S. Woods, Julia M. George, and Chad M. Rienstra.

Mutant protein a30p $\alpha$-synuclein adopts wild-type fibril structure, despite slower fibrillation kinetics. *Journal of Biological Chemistry*, 287(14):11526–11532, March 2012. doi: 10.1074/jbc.M111.306902.

[21] William Humphrey, Andrew Dalke, and Klaus Schulten. VMD – Visual Molecular Dynamics. *Journal of Molecular Graphics*, 14:33–38, 1996.

[22] John Stone. *An Efficient Library for Parallel Ray Tracing and Animation.* Master's thesis, Computer Science Department, University of Missouri-Rolla, April 1998.

[23] Lewis Moffat and David T. Jones. Increasing the accuracy of single sequence prediction methods using a deep semi-supervised learning framework. *Bioinformatics*, 37(21):3744–3751, NOV 1 2021. ISSN 1367-4803. doi: 10.1093/bioinformatics/btab491.
